# Sorghum Root Epigenetic Landscape During Limiting Phosphorus Conditions

**DOI:** 10.1101/2021.05.25.445633

**Authors:** Nicholas Gladman, Barbara Hufnagel, Michael Regulski, Zhigang Liu, Xiaofei Wang, Kapeel Chougule, Leon Kochian, Jurandir Magalhaes, Doreen Ware

## Abstract

Efficient acquisition and use of available phosphorus from the soil is crucial for plant growth, development, and yield. With an ever-increasing acreage of croplands with suboptimal available soil phosphorus, genetic improvement of sorghum germplasm for enhanced phosphorus acquisition from soil is crucial to increasing agricultural output and reducing inputs, while confronted with a growing world population and uncertain climate. Sorghum bicolor is a globally important commodity for food, fodder, and forage. Known for robust tolerance to heat, drought, and other abiotic stresses, its capacity for optimal phosphorus use efficiency (PUE) is still being investigated for optimized root system architectures (RSA). Whilst a few RSA-influencing genes have been identified in sorghum and other grasses, the epigenetic impact on expression and tissue-specific activation of candidate PUE genes remains elusive. Here, we present transcriptomic, epigenetic, and regulatory network profiling of RSA modulation in the BTx623 sorghum background in response to limiting phosphorus (LP) conditions. We show that during LP, sorghum RSA is remodeled to increase root length and surface area, likely enhancing its ability to acquire P. Global DNA 5-methylcytosine and H3K4 and H3K27 trimethylation levels decrease in response to LP, while H3K4me3 peaks and DNA hypomethylated regions contain recognition motifs of numerous developmental and nutrient responsive transcription factors that display disparate expression patterns between different root tissues (primary root apex, elongation zone, and lateral root apex). Suggesting that epigenetic shifts during growth on LP results in targeted gene expression in a tissue-specific manner that optimizes the RSA for improved P uptake.

**Summary:** In response to low P, epigenetic and transcriptional changes stimulate lateral root growth in Sorghum bicolor BTx623, increasing the root surface area for enhanced “mining” of P from the soil.

## Introduction

Sorghum [*Sorghum bicolor* (*L*.*) Moench*] was domesticated in northern Africa ∼6000 years ago (de Wet and Huckabay, 1967; Dillon et al., 2007). A C_4_ grass crop with notable tolerance to drought, heat, and high-salt conditions, sorghum is a top-10 global crop in terms of acreage use. It also serves as a useful model for crop research due to its completely sequenced compact genome (∼730 Mb) (Paterson et al., 2009; McCormick et al., 2018; Cooper et al., 2019) and similarity to crops with larger genomes where repetitive sequences are more prevalent, such as in maize.

Adequate supply of phosphorus (P), which is the most limiting of the three essential plant macronutrients, with finite reserves worldwide (Cordell and White, 2013), limits many cropping systems throughout the world (Calderón-Vázquez et al., 2009). Over 30% of the global cropland are phosphorus deficient (MacDonald et al., 2011), due mainly to P fixation, which is the tendency of phosphate anions to be tightly bound to the surface of soil clay minerals (López-Arredondo et al., 2014). This has been exacerbated because high quality rock phosphate reserves are localized to and extracted from a small number of countries (Cordell et al., 2009). While steady advances in yield have been made across multiple crop species (http://www.fao.org/faostat/en/#data), more will have to be done on the germplasm side to increase overall cropping system PUE.

In recent years, genetic variation in root system architecture (RSA) has been recognized to play an important role in nutrient acquisition efficiency, especially for phosphorus (Lynch, 2011; Bellini et al., 2014). Sorghum adaptation to low-P conditions in the soil is largely governed by P acquisition efficiency with higher root surface area and reduced root diameter acting in concert to enhance P uptake and grain yield (Hufnagel et al., 2014; Bernardino et al., 2019). In fact, a generalized response to low soil phosphorus (LP) involve alterations in root architecture such as those leading to longer and thinner lateral roots, more shallow lateral root angles, and increased root hair production, which all can act to enhance the plant’s ability to mine this diffusion-limited nutrient from the soil (López-Bucio et al., 2002; Schulze et al., 2006; Péret et al., 2014). While there is both intra- and inter-species variation in RSA that are associated with P efficiency (defined in this study as improved crop yield on low P soils) (Miller et al., 2003; Bayuelo-Jiménez et al., 2011), most grasses can show similar structural responses to LP. The molecular mechanisms underlying morphological changes in the root system likely involve a number of signaling cascades (Thibaud et al., 2010; Hu et al., 2011; Hufnagel et al., 2014; Park et al., 2014; Barros et al., 2020) that can be under epigenetic control (Yong-Villalobos et al., 2015) as well as numerous transcription factors (Devaiah et al., 2007; Zhou et al., 2008). Aside from hormone responses in auxin, cytokinin, ethylene, and jasmonic acid pathways (López-Bucio et al., 2002; Thibaud et al., 2010; Li et al., 2012; Nguyen et al., 2013; Khan et al., 2016), there are also notable changes in the epigenetic landscape of the genome that influences transcription factor binding and subsequent *cis*-regulation of gene expression (Smith et al., 2010; Dai et al., 2012; Nguyen et al., 2013; Deng et al., 2016; Chen et al., 2018).

Genetic variation leads to sorghum adaptation to abiotic stress conditions (Doumbia et al., 1993; Doumbia et al., 1998; Hufnagel et al., 2014; Leiser et al., 2014), and there has been recent progress with evincing the genetic components underlying sorghum adaption to soils with low P availability (Hufnagel et al., 2014; Leiser et al., 2014; Bernardino et al., 2019). Specifically, after the identification of the important serine/threonine receptor kinase PSTOL1 in rice (Gamuyao et al., 2012), sorghum PSTOL1 homologs were identified and shown to enhance P acquisition and grain yield under low-P conditions (Hufnagel et al., 2014).

However, much of the epigenetic contribution to nutrient response regulatory networks remains nebulous. Some advancements have been made into regulatory network construction incorporating transcriptomic and regulatory epigenetic regions, such as enhancers, in the related crops of rice and maize (Shi et al., 2014; West et al., 2014; Deng et al., 2016; Zhou et al., 2020), along with investigations of other genes that play roles in plant response to a low phosphorus environment which possibly could be involved in P efficiency (Li et al., 2012; Mora-Macías et al., 2017).

## Results

### Sorghum Root System Responses to Low vs Sufficient Phosphorus Supply

Figure 1 shows representative root images of BTx623 plants grown in our 2D root imaging system. As described in detail in the Materials and Methods, these individual growth systems contain the roots in acrylic chambers, which allow for unrestricted growth of the root systems while maintaining their 2-dimensional RSA and allow for high resolution imaging and analysis of the individual root systems at high throughput. The plants depicted in Figure 1 are representative of the growth response for 5 replicates grown under sufficient P (SP; 200 mM phosphate) or low P (LP; 2.5 mM phosphate), which were imaged at 7, 10 and 14 days after transfer to the SP and LP nutrient solution. Several results are highlighted here: 1) Although sorghum seedlings can produce several seminal roots from the sorghum embryo which later can produce lateral roots, often we see as in the case in Figure 1 that one seminal root dominates and all of the lateral roots emerge from that seminal root. 2) In response to growth on LP nutrient solution, a significant enhancement in lateral root growth is observed.

**Figure 1.**
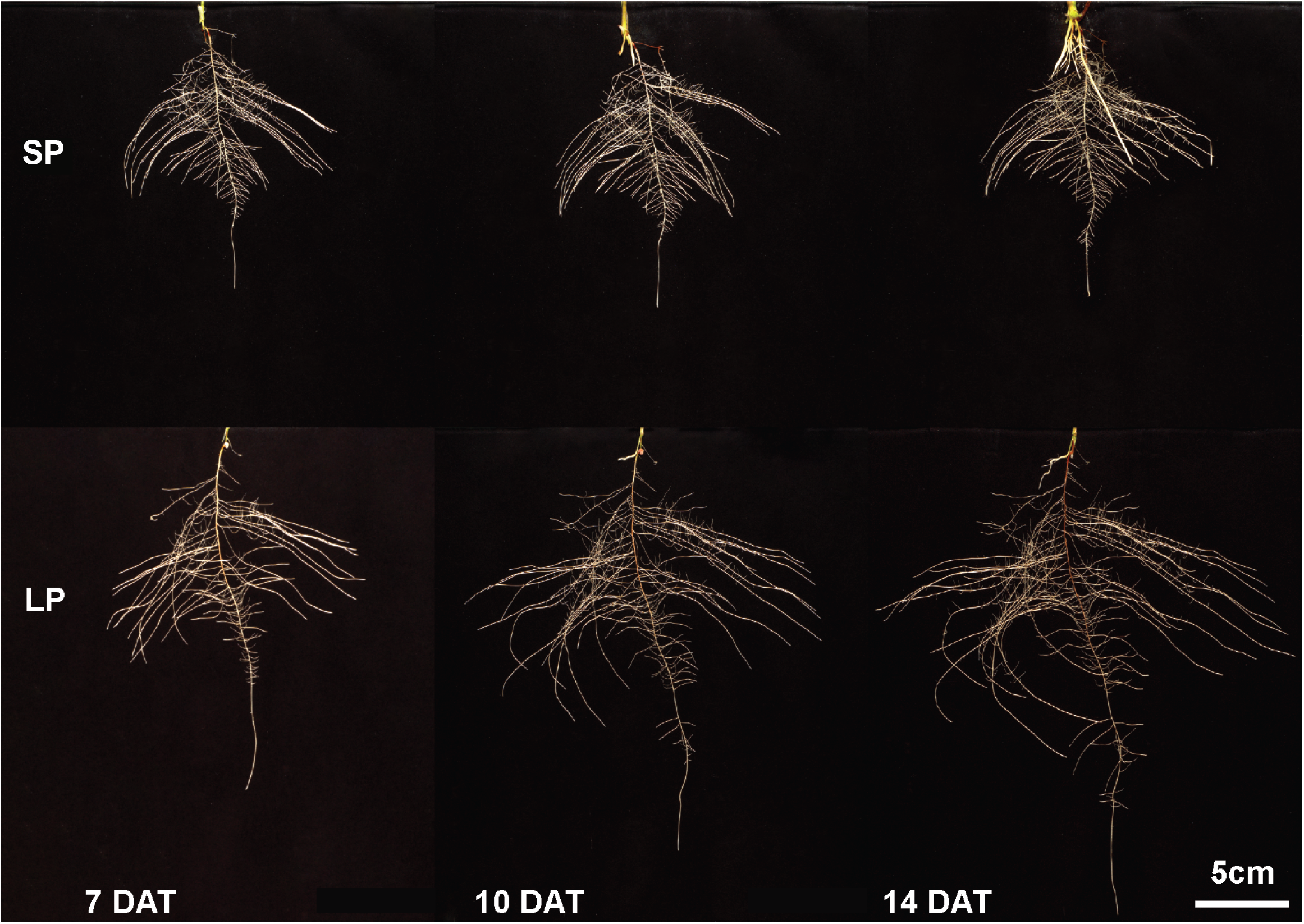
Sorghum Root System Architecture (RSA) under sufficient (SP) and limiting phosphorus (LP). BTx623 seedlings were germinated for 4 days then transferred to hydroponic solutions containing either SP (200 μM) or LP (2.5 μM) nutrient media and grown for an additional 7, 10, or 14 days. Compared to SP conditions, LP stimulates more lateralization of the RSA in addition to deeper root growth through elongation of the apex and intermediate root zones.

The root trait data for BTx623 root systems grown on SP and LP nutrient solution for 7, 10 and 14 DAT are presented in Table 1. It is clear that LP conditions trigger a general increase in both root surface and volume, favoring proliferation of finer roots. The most dramatic low P-induced responses were increases in total root length and root volume (an approximation of root biomass) which is dominated by increases in lateral root growth. A statistically significant decrease in root diameter is also seen by 10 and 14 DAT of growth in LP vs. SP conditions, consistent with the tendency observed at 7 DAT. All of these trait changes are consistent with known alterations in the root system triggered by low P conditions to more effectively explore a greater soil volume and thus more effectively acquire phosphate anions from soils with low P availability.

**Table 1.**
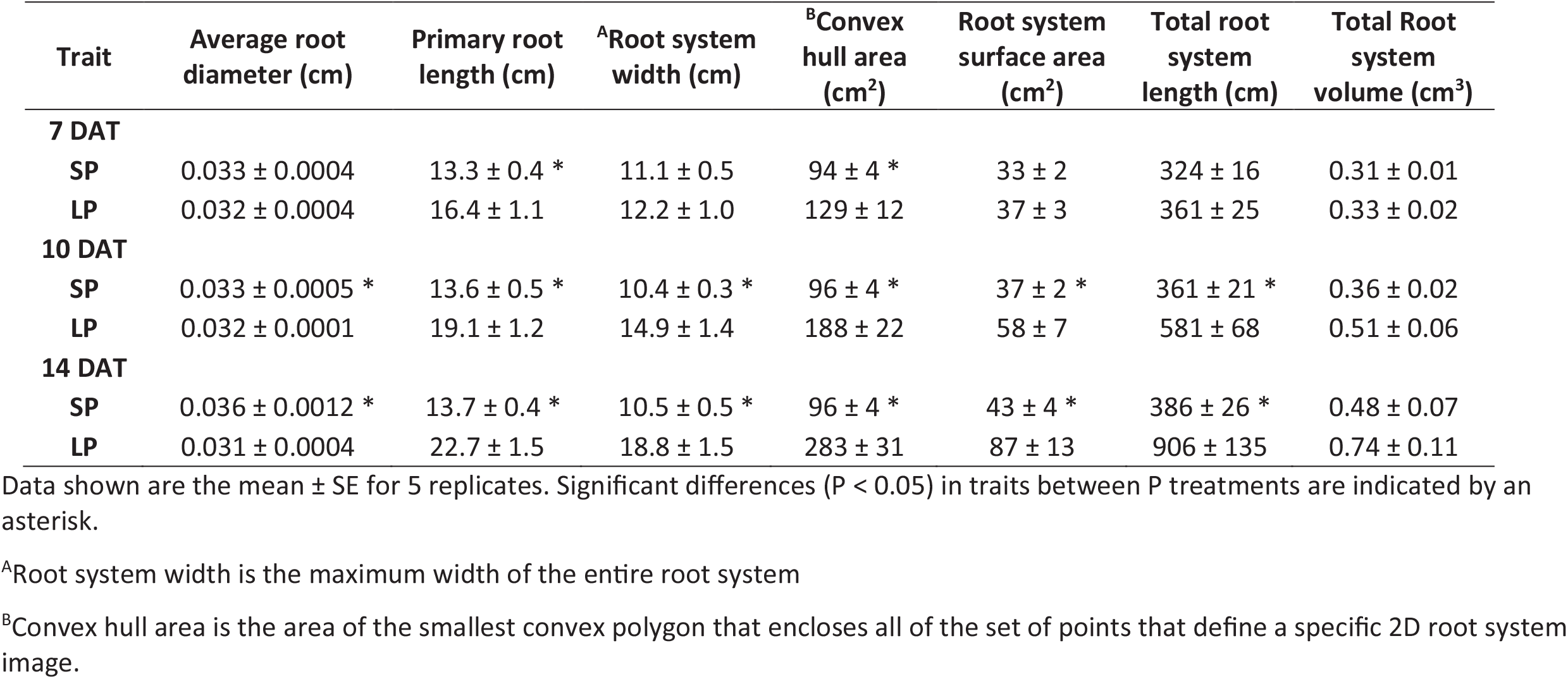
Root system growth and architecture traits quantified for BTx623 grown in nutrient media for 7, 10 and 14 days after transplanting (DAT) under sufficient phosphorus (SP: 200 mM) and low phosphorus (LP: 2.5 mM) conditions

| Trait | Average root diameter (cm) | Primary root length (cm) | <sup>A</sup>Root system width (cm) | <sup>B</sup>Convex hull area (cm<sup>2</sup>) | Root system surface area (cm<sup>2</sup>) | Total root system length (cm) | Total Root system volume (cm<sup>3</sup>) |
| --- | --- | --- | --- | --- | --- | --- | --- |
| **7 DAT** |  |  |  |  |  |  |  |
| **SP** | 0.033 ± 0.0004 | 13.3 ± 0.4 * | 11.1 ± 0.5 | 94 ± 4 * | 33 ± 2 | 324 ± 16 | 0.31 ± 0.01 |
| **LP** | 0.032 ± 0.0004 | 16.4 ± 1.1 | 12.2 ± 1.0 | 129 ± 12 | 37 ± 3 | 361 ± 25 | 0.33 ± 0.02 |
| **10 DAT** |  |  |  |  |  |  |  |
| **SP** | 0.033 ± 0.0005 * | 13.6 ± 0.5 * | 10.4 ± 0.3 * | 96 ± 4 * | 37 ± 2 * | 361 ± 21 * | 0.36 ± 0.02 |
| **LP** | 0.032 ± 0.0001 | 19.1 ± 1.2 | 14.9 ± 1.4 | 188 ± 22 | 58 ± 7 | 581 ± 68 | 0.51 ± 0.06 |
| **14 DAT** |  |  |  |  |  |  |  |
| **SP** | 0.036 ± 0.0012 * | 13.7 ± 0.4 * | 10.5 ± 0.5 * | 96 ± 4 * | 43 ± 4 * | 386 ± 26 * | 0.48 ± 0.07 |
| **LP** | 0.031 ± 0.0004 | 22.7 ± 1.5 | 18.8 ± 1.5 | 283 ± 31 | 87 ± 13 | 906 ± 135 | 0.74 ± 0.11 |

Data shown are the mean ± SE for 5 replicates. Significant differences ( $P < 0.05$ ) in traits between P treatments are indicated by an asterisk.
<sup>A</sup>Root system width is the maximum width of the entire root system
<sup>B</sup>Convex hull area is the area of the smallest convex polygon that encloses all of the set of points that define a specific 2D root system image.

### Root System Transcriptional Responses to Low Phosphorus

Transcriptomic analysis of the seminal primary root apex (first 2 cm), the seminal primary root elongation zone (2-4 cm from root tip), and the root apices of the lateral roots (first 2 cm) reveals that the low P induction of of sorghum homologs of phosphorus starvation induced (PSI) genes was greatest in the lateral root apices (Figure 2A). Based on a gene ontology (GO) analysis within the lateral root apex, genes that displayed an expression change of 2-fold or greater in low phosphorus-grown plants were associated with a number of processes, including different P-related metabolic and signaling pathways, protein phosphorylation, plant response to changes in mineral nutrient availability, lipid modification, and metabolism (Table 2). Other general P-responsive changes in gene expression in the three root regions studied included increased decreased expression of genes involved in metabolic processes, as well as post-transcriptional and translational regulation within the seminal root apex. Also observed was increased expression of genes involved with synthesis of terpenoids and isoprenoids and repression of genes involved in RNA processing and modification, translation and regulation of gene expression in the seminal root subapical region (root elongation zone).

**Table 2.** Gene Ontology enrichment of up-regulated genes with 2-fold expression or higher during LP conditions. Fold enrichment = log_2_ gene expression on LP / gene expression on SP Seminal Root Apex is the first 2 cm of the primary seminal root Seminal Subapical Root Region is the region located 2-4 cm behind the primary seminal root tip Lateral Root Apex is the first 2 cm of the lateral root

Enriched GO Terms in Seminal Root Apex in Response to LP
| GO biological process complete | Fold Enrichment | Raw P-value | FDR |
| --- | --- | --- | --- |
| posttranscriptional regulation of gene expression (GO:0010608) | 0.38 | 8.85E-06 | 6.19E-03 |
| positive regulation of protein metabolic process (GO:0051247) | 0.38 | 8.70E-05 | 4.73E-02 |
| positive regulation of cellular protein metabolic process (GO:0032270) | 0.36 | 6.78E-05 | 4.15E-02 |
| regulation of translation (GO:0006417) | 0.23 | 7.03E-08 | 5.74E-05 |
| regulation of cellular amide metabolic process (GO:0034248) | 0.23 | 4.99E-08 | 4.89E-05 |
| positive regulation of translation (GO:0045727) | 0.04 | 1.54E-09 | 2.51E-06 |
| positive regulation of cellular amide metabolic process (GO:0034250) | 0.04 | 1.54E-09 | 1.89E-06 |
| positive regulation of cytoplasmic translation (GO:2000767) | < 0.01 | 1.04E-09 | 5.09E-06 |
| regulation of cytoplasmic translation (GO:2000765) | < 0.01 | 1.04E-09 | 2.54E-06 |

Enriched GO Terms in Seminal Subapical Root Region in Response to LP
| GO biological process complete | Fold Enrichment | Raw P-value | FDR |
| --- | --- | --- | --- |
| terpenoid biosynthetic process (GO:0016114) | 2.04 | 3.23E-04 | 2.08E-02 |
| terpenoid metabolic process (GO:0006721) | 1.9 | 5.83E-04 | 3.52E-02 |
| isoprenoid biosynthetic process (GO:0008299) | 1.86 | 4.07E-04 | 2.55E-02 |
| isoprenoid metabolic process (GO:0006720) | 1.78 | 2.79E-04 | 1.87E-02 |
| lipid metabolic process (GO:0006629) | 1.3 | 3.17E-04 | 2.07E-02 |
| oxidation-reduction process (GO:0055114) | 1.29 | 1.80E-07 | 2.52E-05 |

Enriched GO Terms in Lateral Root Apex In Response to LP
| GO biological process complete | Fold Enrichment | Raw P-value | FDR |
| --- | --- | --- | --- |
| response to nutrient levels (GO:0031667) | 2.06 | 7.87E-04 | 4.94E-02 |
| phosphatidylinositol metabolic process (GO:0046488) | 1.92 | 4.69E-04 | 3.02E-02 |
| glycerophospholipid metabolic process (GO:0006650) | 1.8 | 1.56E-04 | 1.14E-02 |
| lipid modification (GO:0030258) | 1.78 | 7.56E-04 | 4.81E-02 |
| small molecule catabolic process (GO:0044282) | 1.71 | 1.11E-04 | 8.95E-03 |
| DNA repair (GO:0006281) | 1.68 | 1.00E-06 | 1.14E-04 |
| nucleic acid phosphodiester bond hydrolysis (GO:0090305) | 1.49 | 4.57E-04 | 2.98E-02 |
| chromosome organization (GO:0051276) | 1.45 | 8.70E-05 | 7.22E-03 |
| RNA processing (GO:0006396) | 1.33 | 1.23E-04 | 9.45E-03 |
| phosphorus metabolic process (GO:0006793) | 1.32 | 3.14E-12 | 7.68E-10 |
| protein phosphorylation (GO:0006468) | 1.29 | 2.90E-06 | 3.09E-04 |
| aromatic compound biosynthetic process (GO:0019438) | 1.29 | 4.18E-04 | 2.80E-02 |
| organic cyclic compound biosynthetic process (GO:1901362) | 1.28 | 3.90E-04 | 2.73E-02 |
| cellular nitrogen compound metabolic process (GO:0034641) | 1.28 | 1.47E-11 | 3.28E-09 |
| organic substance biosynthetic process (GO:1901576) | 1.17 | 7.94E-05 | 6.70E-03 |

**Figure 2.**
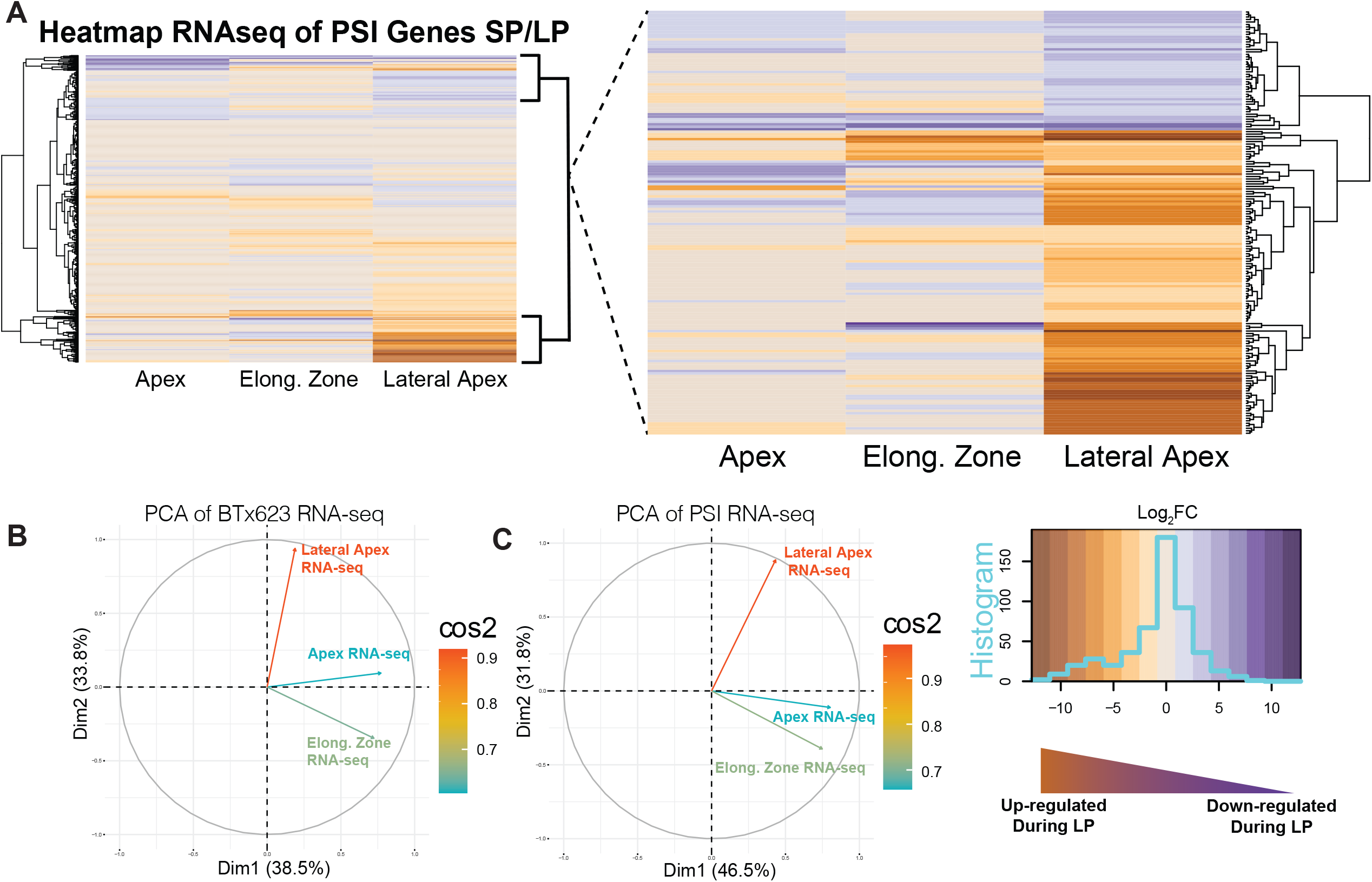
**A)** Heatmap of all sorghum PSI gene orthologs with their root tissue RNA-seq profiles (smaller inset) and an expanded selection of highly differentially expressed PSI genes that shows that most of these genes are concentrated in the lateral root apex. The heatmap legend (lower left of panel) displays the frequency distribution (histogram - cyan trace) of expression of individual genes across the entire range of log_2_ fold change expression values. See Supplemental Data File 1 for topmost up- and down-regulated PSI genes. **B**) PCA analysis of gene expression data with eigenvectors for different root regions, and **C**) PCA analysis showing contribution of all PSI genes; importance of contribution is displayed as cos2.

Upon inspecting distinct genes families, several well-known PSI genes were strongly upregulated in response to low P in the lateral root apices (Supplemental Data File 1 and 2). Many of these PSI transcripts belong to transporters, metabolic, and transcription factor family genes, and for these genes, in general their largest change in expression in response to low P was in the lateral root apex. To determine if a particular root region or gene family was responsible for the greatest variation within the transcriptomic data, a principal component analysis (PCA) was performed against the RNA-seq expression data for all BTx623 genes as well as for specific gene families (families associated with PSI, metabolic pathways, transcription factor families, *etc*.). No specific gene family was found to be clustered in the PCA, but gene expression in the lateral root apex displayed a noticeable difference in correlation compared to either the root apex or root elongation zone based on their PCA eigenvectors direction and intensity (Figure 2B, C). This indicates a stronger transcriptional correlation between the root apex and subapical regions versus the lateral root regions with regards to low P induced changes in gene expression.

When evaluating subsets of other notable gene families, such as transcription factors and specifically the DNA binding domains of TF families involved in plant stress responses, such as AP2, WRKY, NAC, bZIP and B3, the lateral root apex displays a larger number of genes that also exhibit a greater up-regulation of P deficiency response genes compared to either the seminal root apex and the seminal root sub-apical regions. The later regions display either negligible or a greater number of P responsive down-regulated genes. A gene ontology (GO) analysis of these transcription factor gene families that are strongly up-regulated (2-fold expression or greater) in the lateral root regions shows an enrichment of auxin-activated signaling pathway genes (GO:0009734), whereas the down-regulated genes in the root apex are enriched for response to chitin (GO:0010200) and abscisic acid (ABA)-activated signaling pathway (GO:0009738), as well as the auxin-activated signaling pathway (GO:0009734). The strongly down-regulated WRKY transcription factors that are a part of the canonical PSI genes are paralogs involved in the salicylic acid pathway and had association with fungal and chitin-responsive pathways (SORBI_3002G202800 and SORBI_3002G202700), displaying potential overlap of biotic and abiotic stress responsive pathways.

### DNA and Histone Methylation Profile Throughout the Sorghum Genome

ChIP-seq profiling of Histone 3 (H3) lysine trimethylation showed distinctive signaling at different regions throughout the sorghum genome in the intergenic and genic space. H3K27me3 (repressive) and H3K4me3 (activation) marks had similar averaged global coverage nearby and within genic regions (Figure 3A). However, there was a drop (∼ 50%) in H3K4me3 enrichment within the TSS, genic, and 5’ prime UTR regions under LP conditions compared to SP. When interrogating intergenic regions, H3K4me3 enrichment increased by ∼50% in response to LP growth conditions compared to SP, whereas K27me3 enrichment remained mostly unchanged under either phosphorus growth treatment in the intergenic space (Figure 3B). Clustering analysis revealed that H3K4 trimethylation can be grouped into three primary Kmeans clusters: upstream, from >3000 bp to the transcriptional start site (TSS), genic, and downstream/non-specific (Figure 3C, D). The H3K27 trimethylation Kmeans clusters cover genic, downstream (from transcriptional end site (TES) to >-3000 bp), and upstream/non-specific regions. In agreement with the enrichment observations, the heatmap analysis indicates that both the H3K4me3 and H3K27me3 signals appear to decrease across all regions under low phosphorus, with K4me3 decreasing noticeably more than K27me3. The heatmap analysis stops at 3 Kb upstream or downstream of the TSS and TES, respectively. So, the increased enrichment of the H3K4me3 in intergenic spaces under LP conditions (Fig 3A) must originate from further away than these margins, suggesting a role of enhancer elements or more distant promoter-binding locations.

**Figure 3.**
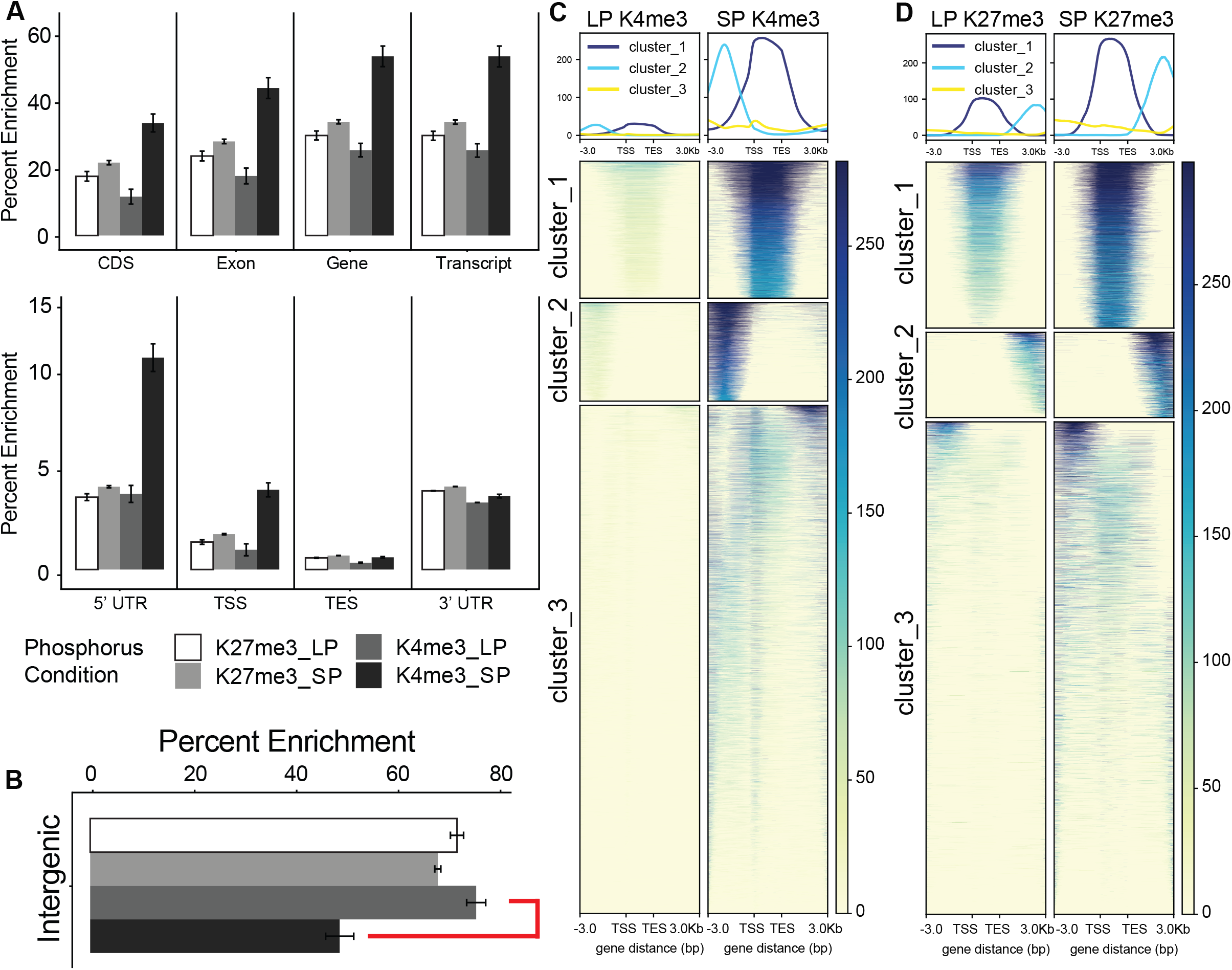
Epigenetic Profiles of BTx623 whole root systems under normal and limiting phosphorus. **A**) Global enrichment profiles of sorghum H3K27me3 and H3K4me3 across gene model components. **B**) Global enrichment profile of K4me3 and K27me3 epigenetic marks across intergenic space during LP and SP. **C**) Global heat map and profile graphs of H3K27 and H3K4 trimethylation around gene models. K-means analysis generated 3 major clusters that roughly segregate methylation occurrences to genic, upstream, and/or downstream regions during normal and LP conditions.

Sequence motif analysis revealed both shared and discrete DNA-binding motifs for both K27 and K4 histone methylation peaks, with certain motifs specific for growth on SP conditions and others specific for LP conditions (Figure 4). The recognition sequence for the APETALA2/ethylene-responsive element binding protein (AP2/EREB) transcription factors was enriched in both K4me3 and K27me3 peaks. While the DNA-binding sequence recognized by AP2/EREB family proteins was detected, the exact position weight matrices between P conditions varied. This could suggest that either a subset of AP2/EREBs are being regulated in response to P starvation based on their precise *cis*-binding motif or it could be reflective of a broad recognition profile for the AP2/EREB transcription factors. Other multiple transcription factor recognition-binding motifs were present upon inspection of the K4 and K27 methylation footprints. These included, but were not limited to, Trihelix, WRKY, NLP, NAC, ABI3, BZIP, and B3-domain containing TF families.

**Figure 4.**
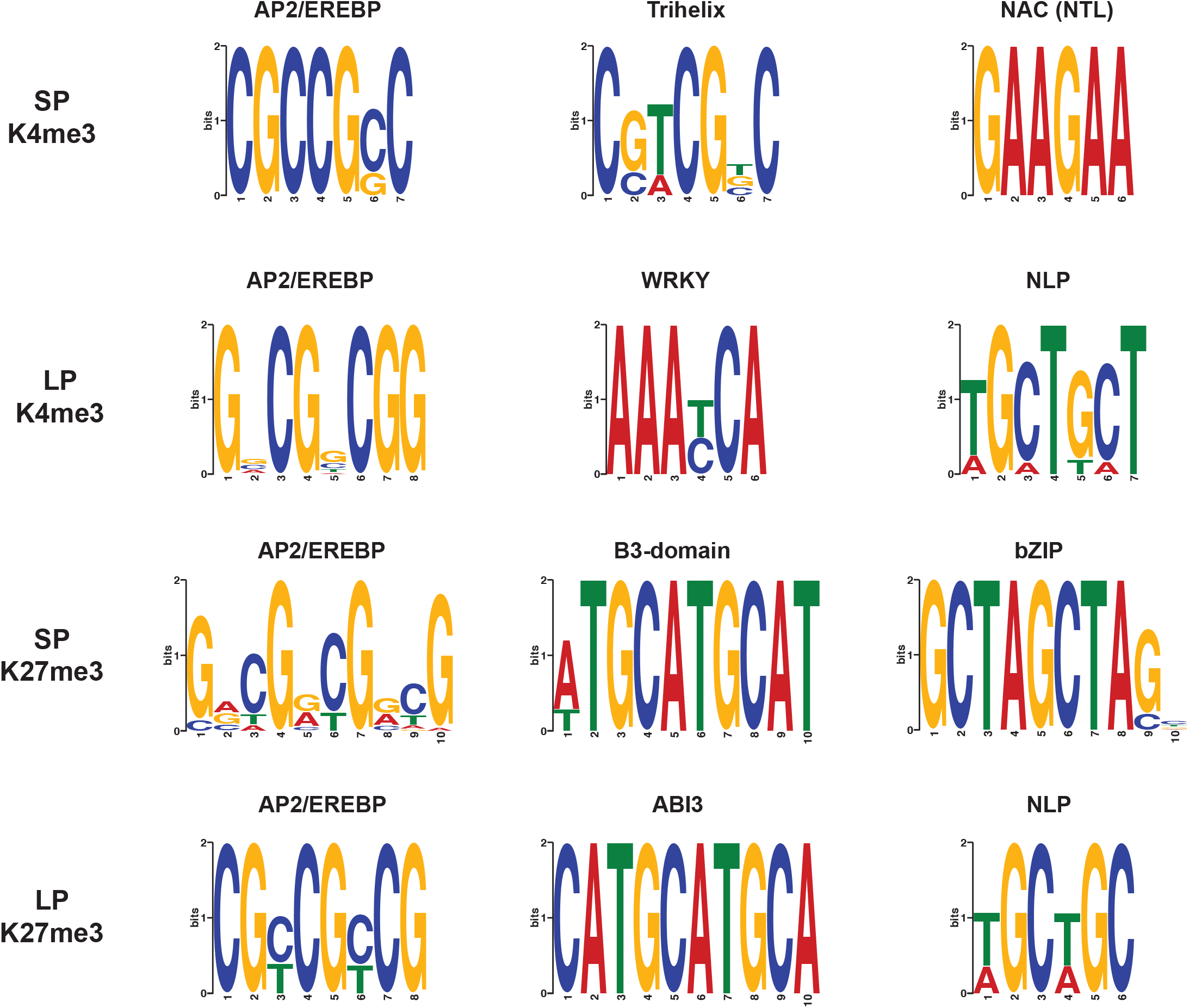
DNA Binding Motif Enrichment Analysis. Global MEME motif analysis of the 100 bp flanking sequences from K27me3 and K4me3 peaks during SP and LP conditions. All motifs were detected using the ANR method in the MEME suite program with minimum significance threshold of e < 0.01 or less.

Global DNA methylation (5-methylcytosine) enrichment levels remained mostly unchanged from SP to LP conditions across averaged genic or intergenic regions (Figure 5A, B) and decreased globally over genic regions, in line with previous observations for actively transcribed loci. DNA methylation levels were enriched the most in intergenic and transcribed regions and the least at the start and stop codon positions and at the three and five prime UTRs. DNA methylation was lower in exon regions compared to entire genes and transcripts, indicating increased methylation at the intron and non-coding gene space. Differential CG methylation in response to growth on LP conditions was the predominant form of differential methylated regions (DMRs) and was more common at surrounding gene elements (promoters, exons, and introns) than either CHG or CHH methylation (Figure 5C and Figure 6). Most of the CHG and CHH differential methylation (70-80%) was localized to the intergenic space (Figure 6B, C). CG hypomethylation is more common across the genome and specifically at genic and promoter regions; hypomethylation is also the most common form of CHG DMR, but such changes were fairly similar at genic and promoter regions when compared to hypermethylation (Figure 6B, C). Notably, almost 100% of the statistically detected differential CHH methylation was hypomethylation. Motif analysis of regions surrounding differentially methylated CG marks were found to be enriched for AP2/EREB, WRKY, NLP, TCP, bZIP, C2H2, BZR, trihelix, and NAC family transcription factor binding sites (Supplemental Figure 1). Additionally, CHG marks were enriched for AP2/EREB, WRKY, NLP, TCP, E2F-like, and LOB (Lateral Organ Boundary) transcription factors (Supplemental Figure 2). DNA surrounding hypomethylated CHH loci were enriched for AP2/EREB, WRKY, NLP, TCP, and NAC transcription factor binding sites, amongst others (Supplemental Figure 3).

**Figure 5.**
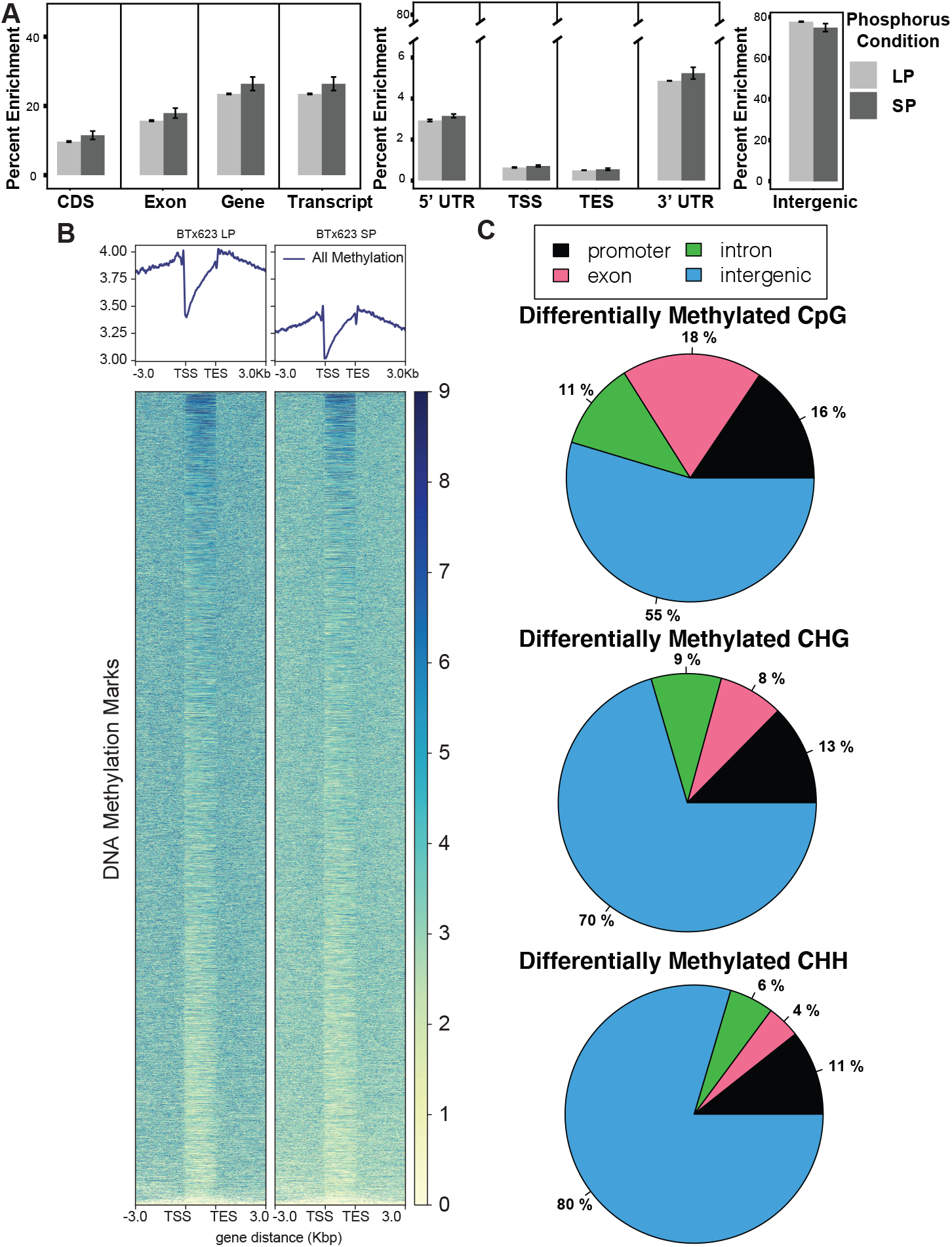
BTx623 Global DNA Methylation Profile. **A**) Global enrichment profiles of DNA methylation (BS = CG, CHG, and CHH) across gene model components during LP and SP conditions. **B**) Heat map of 5-methylcytosine methylation across gene models in limiting (LP) and sufficient (SP) phosphorus. **C**) Differential DNA methylation (hyper- or hypo-methylated) changes over promoter, exon, intron, and intergenic sequences in LP conditions compared to SP. Almost 100% of differential CHH methylation is hypomethylation.

**Figure 6.**
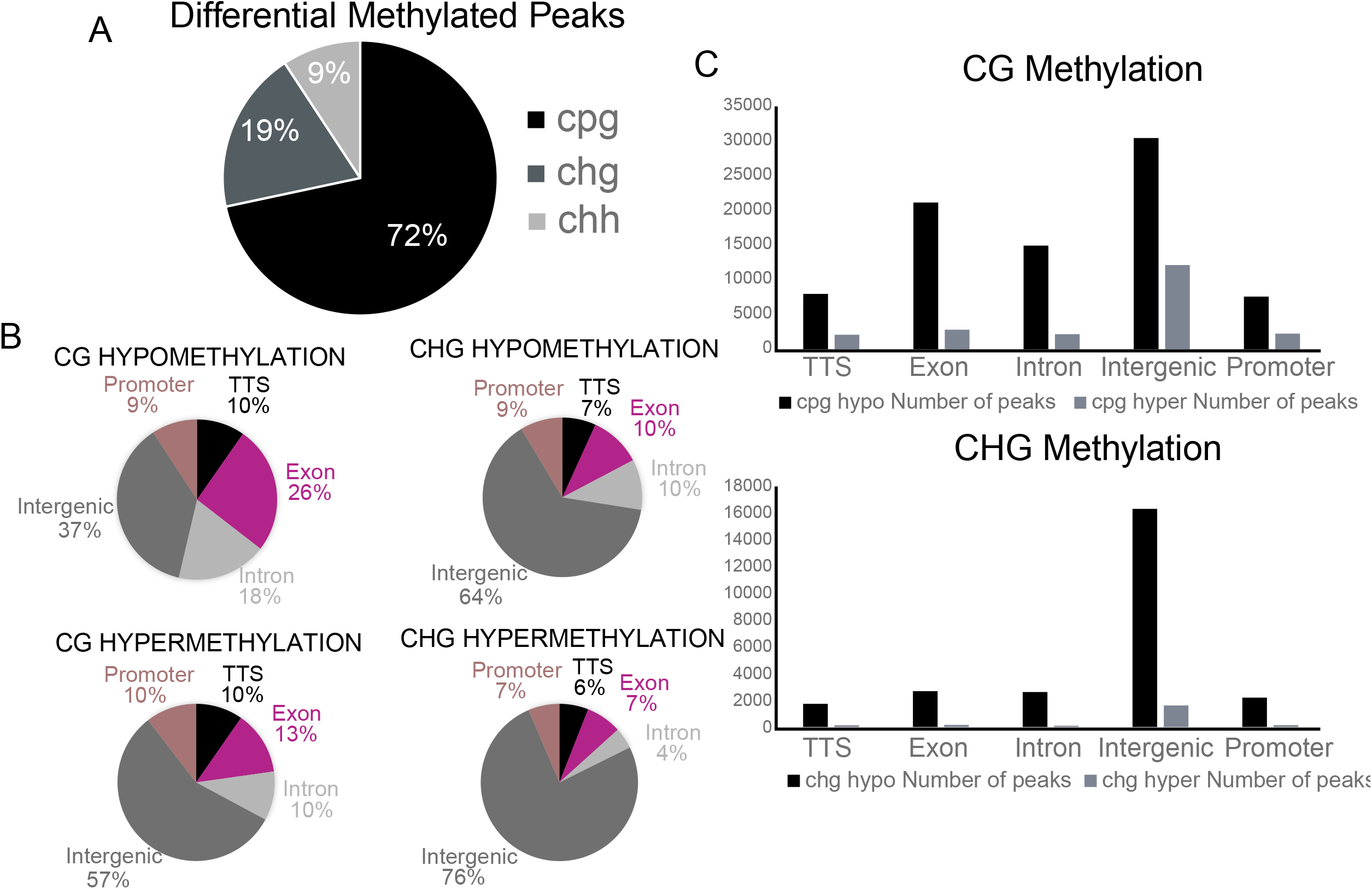
Differential DNA Methylation Regions in BTx623. **A**) Breakdown of all differentially methylated peaks (regions) by CG, CHG, and CHH in the BTx623 genome under LP growth conditions compared to SP growth. **B**) Gene model breakdown of CG and CHG hyper- and hypo-methylation in LP grown plants compared to SP. CHH DMRs are ∼100% hypomethylated in roots of LP-grown plants. **C**) Total number of CG and CHG DMRs by hyper- and hyper-methylation by genomic location.

The global change in histone trimethylation can also be observed by taking the ratio of K4 and K27 marks in the genic, 1000 and 5000 bp upstream bins and seeing how their frequency distribution changes (Supplemental Figure 4). Across all intervals during growth on LP, the K4/K27 ratio distribution shrinks to a narrower range of values (specifically a lower ratio of K4 to K27) when compared to SP, suggesting that one or both marks are becoming less common and/or more consolidated across the epigenome in response to P limiting conditions.

Upon analysis of selected transcription factor gene families identified from the MEME analysis above and presented in Figure 5, it was determined that subgroups within these gene families display strong up- or down-regulation of expression in response to LP conditions (Supplemental Figure 5). Specifically, a number of NAC, B3-domain containing, bZIP, WKRY, and AP2 transcription factors showed strong upregulation in the lateral root apices, and to a lesser extent in the primary root regions.

### Phosphorus Responsive Regulatory Networks

To visualize complex interactions and contributions during root responses to LP growth conditions, protein-protein interaction networks were generated using the STRING database. The gene nodes were then filtered to reduce the network down to those genes that had associated K4me3 peaks; the gene nodes were then colorized to correspond to their expression at lateral root regions. This phosphate responsive root gene regulatory network (GRN) was organized in a hierarchical radial layout to better visualize both transcription factor interactions and biosynthetic and metabolic pathways (Supplemental Data File 3). Screening for interacting gene products that are up- or down-regulated during low P conditions revealed several distinct families. Numerous auxin, autophagy, and photosynthetic genes show distinct regulatory patterns within the GRN, as do several genes involved in sulfur and cysteine metabolism. Two up-regulated cysteine-related metabolism and signaling genes in particular were SORBI_3008G179900 and SORBI_3001G427300, which encode an O-acetylserine (thiol) lyase (OAS-TL) and a glycosyl transferase, respectively. Prior research into hypomorphs of these orthologs have shown they are either associated with reduced root growth and root surface area, specifically within lateralization of roots or are essential for proper response to LP conditions(Okazaki et al., 2013; Wang et al., 2015; Mei et al., 2019). Upon examination of the enriched peaks of H3K4me3 and H3K27me3 sequencing across the OAS-TL and transferase gene models, there is a noticeable enrichment for H3K4me3 peaks at the five prime genic or promoter region in both genes in LP or SP-grown plants (Supplemental Figure 6), but there is a narrowing of the peak width under LP growth conditions.

## Discussion

### BTx623 Root Responses to P Deficiency Stress

We observed that P starvation caused a rather drastic remodeling of root system morphology in BTx623. From 7 DAT to 14 DAT, under low P growth conditions, root system length, surface area and volume increased 250%, 235%, and 224% respectively, compared to the same root traits for plants grown on sufficient P (Table 1). Furthermore, at 7 DAT there was no difference in average root diameter between the two P treatments, but by 14 DAT, root system diameter had decreased a moderate (14%) and statistically significant amount in LP *versus* SP grown plants. Increases in lateral root growth in response to P deficiency have been previously observed in other plants, for example, in beans (Lynch and Brown, 2001), Arabidopsis (Arrese-Igor et al., 2020), and tomato (Hai-Bo et al., 2001). These changes in root traits are consistent with P acquisition efficiency explaining most of the genetic variation in overall P efficiency in maize and sorghum (Parentoni and Souza Júnior, 2008; Parentoni et al., 2010; Morais de Sousa et al., 2012; Bernardino et al., 2019). Phosphorus uptake is hindered, in part, due to the phosphate anion interacting strongly with Al-and Fe-oxides on the surface of soil clay minerals, resulting in much of it being fixed in the soil. This chemical behavior between P and soil minerals results in P being the most diffusion limited of the essential mineral nutrients. Hence plant developmental strategies that increase root proliferation in the soil to reduce the distance between roots and soil P in ways that reduce the carbon cost for producing more roots, such as generating finer roots that have a higher surface area to volume ratio, appear to be key aspects of P efficiency. Certainly, the root responses shown here for sorghum BTx623 are consistent with these strategies.

### Gene Targets for Crop P Efficiency Improvement

In a broad sense, the root transcriptome data presented in this study is consistent with prior investigations on the genes whose expression is significantly altered by P deficiency stress and are one resource for candidate P efficiency genes. These include transcription factors involved in root development, such as *SHORTROOT* (*SHR*), *SCARECROW* (*SCR*) and *ROOTLESS CONCERNING CROWN AND SEMINAL ROOTS (RTCS)*, phosphate transporters, and genes associated with auxin signaling, response, and transport. The significant changes in gene expression induced by P deficiency primarily in lateral root apices (compared to the primary root apical and subapical regions) depicted in Figure 2 correlate at the physiological level with the root system response to LP in Figure 1 and Table 1, where lateral root growth stimulation is the major P deficiency growth response.

When evaluating transcription factors that are strongly up- or down-regulated in the lateral and apical root regions, a biological process ontology analysis suggests that specific hormone shunting is occurring throughout the RSA, with a specific increase in auxin activation within the lateral roots, and a shared downregulation of auxin and ABA signaling possibly occurring in the root apex. There could also be an overlapping response that is generally believed to be a response to biotic stress but could also be involved in root low P responses. This biotic response that plays a role in modulating RSA in response to chitin, involves the downregulation of WRKY transcription factors (Supplemental Data File 1) that play roles in root responses to fungal chitins that alter root system architecture (Wang et al., 2019; Wang et al., 2020).

An interesting finding from the gene expression and histone marks data was the identification of multiple genes involved in sulfur and cysteine metabolism that were discovered through the generation of a lateral root gene regulatory network (GRN) filtered for H3K4me3 peak enrichment and up-regulated expression in response to LP. The functional relevance of this finding is supported by previous work indicating that sulfur metabolic pathways are integral to auxin changes in the root apex, leading to enhanced lateral root growth (Romero et al., 2014; Zhao et al., 2014; Li et al., 2015; Wang et al., 2015; Mei et al., 2019). It is not surprising to find such links between regulatory networks for two different essential nutrients, in this case, P and S. Another example of such interactions comes from the growing evidence of cross talk and co-regulation between P and N homeostasis (Medici et al., 2019) provide evidence that the plant P starvation response is regulated in part by both local and long-distance N signaling pathways. Another fascinating regulatory interaction of essential mineral nutrients is the complex interplay and role of Fe transport, cellular compartmentation and metabolism in P deficiency-induced alterations in primary root growth (Müller et al., 2015; Balzergue et al., 2017; Mora-Macías et al., 2017).

The DNA and histone methylation footprints presented here agree with prior investigations into epigenetic profiles of genic regions as we found there was significant enrichment at the gene body for both K27me3 and K4me3, with a drop in DNA methylation over the same space (Regulski et al., 2013; Zhu et al., 2017; Peng et al., 2019). Specifically focusing on histone methylation, K4me3 levels in LP plants seemed to decrease significantly across the TSS, five prime UTR, and transcript-encoding components of gene, and while K27me3 levels also decreased across the same segments, it was not nearly to the same degree as K4me3 levels (∼50% drop). Another distinct global histone methylation characteristic in response to LP was the increase in H3K4me3 levels in the intergenic space. This is contrasted with global intergenic H3K27me3 and DNA methylation levels that remained largely unchanged under LP versus SP treatments. Specifically, the spread of the H3K4me3/H3K27me3 ratio across intergenic regions became narrower under limiting phosphorus, more so at loci proximal to transcriptional start sites (>1000 bp upstream). Additionally, it was observed that there are moderately weak positive correlations between gene transcription and H3K4me3 mark intensity at genic and promoter regions and weakly negative or negligible correlations with H3K27me3 mark intensity across the same intervals (data not shown; ∼0.3-0.4 spearman correlation values for transcript CPM compared to K4me3 ChIP-seq peaks and ∼-0.1-3 for H3K27me3 peaks). All of these findings suggest that such histone methylation patterns could play a role in overall phosphate starvation adaptation. Further investigations into the frequency and location of more open chromatin marks, such as H3 lysine acetylation, might reveal stronger correlations with gene expression. Additionally, it was discovered here that enriched sequences surrounding histone methylation sites and DNA hypomethylation regions included binding sequences for transcription factors known to be associated with root growth and response to environmental stimuli (Figure 4). These data set the stage for future investigations into promoter, enhancer, and cryptic sequence regulatory candidates that may play roles in the molecular basis for enhanced P efficiency, which could be used to improve crop adaptation to low P soils which are prevalent worldwide, limiting agricultural production especially in the developing world (Lynch, 2011; Kochian, 2012).

During a shift from growth on sufficient to low phosphorus conditions, there is a coordinated de-methylation in sorghum roots that occurs both at the K4 and K27 lysine in histone 3, along with a trend towards increases in hypomethylated DMRs for 5-methylcytosine DNA methylation. Specifically, K4me3 peaks at the genic and promoter regions have a mild positive correlation with gene expression. The shift in epigenetic regions containing K4me3, K27me3, or DNA methylation peaks in response to low P are enriched for numerous DNA-binding motifs recognized by several transcription factor families. Some of which are notably, AP2/ERF, NAC, WRKY, and LOB transcription factors that can play a role in downstream gene activation for numerous hormone signaling pathways, specifically auxin, as well as canonical PSI genes and shared nutrient response pathways (nitrogen and sulfur). These gene up-regulations seem to be localized predominantly to the lateral root region, suggesting that the primary transcriptomic remodeling of the RSA is focused within the lateral root apices, and that the RSA epigenetic shift likely supports these changes in RSA to maximize root growth in low soil phosphorus environments (Supplemental Figure 7).

## Conclusions

Imposing phosphorus (P) deficiency stress induced significant changes in the root architecture of the Sorghum bicolor reference cultivar, BTx623. These included a significant stimulation of lateral root growth, which resulted in increases in total root system length, root surface area and total root system volume. We used genomic and epigenetic approaches to investigate transcriptional and chromatin changes in response to these P deficiency-induced changes in root architecture, which enhance sorghum’s ability to acquire P from the soil. Three distinct root regions were studied: the primary root apical and subapical regions, and the lateral root apical region. Intriguing transcriptional responses were seen, with the majority of the genes whose expression were significantly up- or down-regulated localized to the lateral root apex. Under LP growth conditions, the epigenetic profiles of DNA methylation and histone trimethylation also showed distinct changes both around gene models and especially in the intergenic space. Combined together, these data revealed unique epigenetic shifts under limiting nutrient conditions that were associated with genes that could influence root development, but also previously undiscovered candidate genes and regulatory motifs that could influence changes in sorghum RSA that enhances P acquisition from low P soils. Taken together, the transcriptomic and epigenetic data in this study are a resource for future research on sorghum and other related cereals for the ultimate purpose of generating improved crop PUE through, in part, root systems that are more efficient and effective at “mining” P from the soil.

## Methods

### Plant Growth Conditions

#### Sorghum growth protocols for epigenetic and genomic assays

Sorghum seeds were surface-sterilized in a solution of 6% (v/v) sodium hypochlorite for 15 minutes. The sodium hypochlorite was removed by washing the seeds with distilled water 5 times. The seeds were then germinated on filter paper moistened with water for 4 days. After that, uniformly growing seedlings were transferred to 200 liter polypropylene tubs grown hydroponically with aerated modified Magnavaca nutrient solution (see (Magnavaca et al., 1987) for nutrient solution composition). For nutrient solution with sufficient P (SP), the phosphate concentration was 45 μM and for low P (LP). The phosphate concentration used was 2.5 μM. The sides of the tub were covered with white light reflective plastic and the nutrient solution in each tub was covered with grey, closed-cell polyethylene foam strips (McMaster-Carr, Elmhurst, IL, USA) to prevent light penetration and support the seedlings during the experiment (Fig. 1b). Plants were grown under controlled conditions in an EGC walk-in growth chamber for the duration of the experiment (27 °C day/22°C night, 12/12 h photoperiod, 500 mmol m-2 s-1 photons). After 1 day in sufficient P nutrient solution, half of the seedlings were transferred to low phosphorus nutrient solution (LP) where the 45 μM KH_2_PO_4_ was replaced with 2.5 ×μM KH_2_PO_4_. The pH of the solution was maintained at the range of 5.6-5.7 by adjustment with 0.5 M NaOH or HCl every 2 days. The nutrient solution was renewed every 5 days. Each treatment had 3 biological replications. Plants were grown in SP and LP conditions and root tissue was harvested after 10 days of growth on LP or HP conditions, for preparation of the libraries described below.

#### Sorghum plant growth for 2d root system architecture (RSA) analysis)

Sorghum seeds were surface sterilized and geminated as described above. After 4 days of germination, uniform sorghum seedlings were transplanted and grown hydroponically using the same modified Magnavaca nutrient solution described above except the phosphate concentration used for SP media was increased to 200 μM. As above, the pH of the solution was maintained at the range of 5.6-5.7 by adjustment with 0.5 M NaOH or HCl every 2 days. The nutrient solution was renewed every 2 days for the growth of the plants in the pouch system. Single sorghum seedlings were grown in specially designed and constructed acrylic chambers which hold the seedling and root system on the surface of filter paper (described in more detail below. These 2D root growth chambers were placed vertically into 100 L polypropylene tubs (L ×W× H= 50 × 30 × 65 cm) containing aerated modified Magnavaca nutrient solution whose composition is given above. Aluminum rails attached to the longer sides of each tub allowed the growth pouches to hang into the nutrient solution in the tubs. The level of the aerated nutrient solution was adjusted so it always was just below the root apex of the fastest growing seminal root. Each container held 20 plants, each in their individual pouches. A cover is placed on top of and sides of the tubs allowing the shoot to grow while excluding light from the roots and the nutrient solution.

The plant growth pouches consist of several layers of different materials. On the bottom is a perforated plastic back that provides rigidity to the pouch and the holes in the plastic backing facilitate better nutrient and air exchange. On top of this plastic back were placed several layers of germination paper which provide the nutrient solution to the plant roots. On top of the germination paper is a single sheet of black filter paper that serves as an anchor for the roots as they grow across its surface, and the black background provides high contrast with the roots, greatly improving root imaging. Finally, a sheet of clear polyester film covers the root system and black filter paper, protecting the root system from drying out and also contributing to root system anchorage to the black filter paper during growth, maintaining the root systems two-dimensional architecture.

### Sorghum Root Imaging and Analysis

At 7, 10 and 14 days after transplanting, individual pouches with one sorghum plant were removed from the hydroponic tub, the plastic film carefully removed, and the plant on the pouch system was carefully placed oriented horizontally in a glass tray containing water with the root system totally submerged in the water. The tray with plant was placed into a 2D root imaging platform consisting of a frame constructed from 80:20 aluminum rails, to which is attached a Nikon D7200 DSLR camera with a 50 mm lens. Also attached to the frame are flashlights with diffusers (soft boxes) to diffuse the illuminating light across the root system, preventing shadows that can compromise the quality of root images. The root systems are submerged in water to prevent any unwanted diffraction and reflection of the illuminating light.

Our Plant Root Imaging and Data Acquisition (PRIDA) program, which is a python-based image acquisition and data management software, was used for collecting and storing raw images and project meta-data into a single Hierarchical Data Format (HDF5) file (www.hdfgroup.org). Images were then extracted as TIFF image files for further processing and root trait computation. These TIFF images were then fed into commercial and publicly available software packages for root trait computation. These software packages process the images to segment the root system architecture and separate it from other background objects by using different techniques, such as global, local, or adaptive thresholding. The segmentation process provides a clean root system architecture and for the next step in the process, skeletonization, which provides a single pixel line to provide a skeleton of the root system architecture. This skeleton is used to estimate a range of root growth and architecture traits. WinRHIZO software (Regents Instruments, Inc) was used to quantify root growth and topology traits, and GiA Roots (Galkovskyi et al, 2012) was used to quantify 2D root architecture traits. Taken together, root topology and architecture traits including average root diameter (cm), primary root length (cm), root system width (cm), convex hull area (cm^2^), specific root length (cm/cm^3^), root system surface area (cm^2^), total root system length (cm) and total root system volume (cm^3^) were investigated in this study.

### RNA-seq Library Preparation and Analysis

The first two cm of the seminal root tips were excised, and then the next 2 cm of the same seminal roots were also excised, and both tissue samples were separately frozen in liquid N_2_. Additionally, the first two cm of the lateral root tips were also excised and frozen, and all three tissue samples were used for preparation of the different libraries described here and below. Root tip samples were collected from both SP and LP grown plants, after 10 days of growth in SP and LP media.

Root tissue samples were ground in liquid nitrogen and then the RNA was isolated in Trizol reagent (ThermoFisher) followed by purification using Zymo-spin columns (Directzol RNA kit; Zymo Research). Column-bound RNA was treated with DNase 1 and finally eluted with 50 uL of DNase/RNase-free water. RNA quality was determined using the Bioanalyzer hardware with the Agilent RNA 6000 Nano Kit (Agilent). Poly-A RNA was isolated from total RNA using Dynabeads (ThermoFisher). Libraries for sequencing were created using the ScriptSeq RNA-seq prep kit (Illumina) following manufacturer protocols. Final libraries were amplified with 17 cycles of PCR and assessed on the Bioanalyzer with the High Sensitivity DNA kit (Agilent). All root tissue libraries that were sequenced comprised two biological replicates.

#### Sequencing platform information

Sequencing was performed at Cold Spring Harbor Laboratories using the Illumina HiSeq2000 platform with 100 bp paired-end reads. Paired-end fastq files were trimmed for quality with Trimmomatic and then merged with Samtools before aligning to the v3.4.1 Sorghum bicolor genome with Kallisto package in rStudio v3.6.1. Differential gene expression was determined through DESeq2 after importing Kallisto abundance files with tximport package. Dimensional analysis was performed using the factoextra package. Heatmaps and hierarchical clustering were performed with the heatmpap.2 package in rStudio.

The phosphate starvation Induced (PSI) gene set was derived from the Yong-Villalobos (2015) manuscript. See Supplemental Data File 2.

### ChIP-seq Library Preparation and Analysis

The entire root systems were harvested for ChIP-seq library preparation. The root tissue was cut into sections and fixed in 10 mM Dimethyl adipimate (DMA; Sigma-Aldrich) while applying a vacuum. Samples were ground in liquid nitrogen and then placed in extraction buffer, filtered through miracloth and then a 30 mm CellTrics filter (CellTrics). Nuclei were isolated and then chromatin was sheared in 130 mL tubes (Covaris). Shearing was performed on a Covaris S220 sonicator for 5 minutes at a cycle/burst of 200, peak power of 175, and duty factor of 10. Immunoprecipitation was performed with antibodies for H3K27me3 (Millipore), H3K4me3 (Abcam), H3 (Abacam) for input control, and Protein A Dynabeads (ThermoFisher) for input control. Crosslinking was reversed and samples were purified using Chip DNA Clean and Concentrator Kit (Zymo Research). Libraries were created using the Nextflex ChIP-Seq library prep kit (Bioo Scientific) with the NextFlex ChIP-seq Barcodes. Sequencing was done on a HiSeq 2500 v4 with 125 bp paired-end reads. Paired-end fastq files were trimmed for quality with Trimmomatic and then mapped to the v3.4.1 Sorghum bicolor genome with BWA (bwa mem -q 30, -f 2 -ubhS). Sorted bam files were marked for duplicates with Picard and then merged with Samtools. Tag directories creation and peak calling, peak enrichment comparison was performed through the Homer package. Overlapped peaks were identified through Bedtools (intersectBed). Peak annotation was performed using the annotatePeaks module of the Homer software. Samtools was used to obtain flagstat metrics and perform various file conversion and handling. Various visualization and file creation was perfomed using the deepTools package (Ramírez et al., 2016); aligned bam files (that were input-controlled) were converted to bigWig format for generating computeMatrix files for plotHeatmap and plotEnrichment outputs. Kmeans cluster number was determined by selecting the result that generated the distinct read coverage within and around gene models while minimizing cluster overlap.

### Bisulfite-seq Library Preparation and Analysis

Bisulfite-sequencing was done as described in (Li et al., 2014). The root apices (first two cm) of all primary and lateral roots were excised from plants grown as described above after 10 days of growth on SP and LP media. DNA was isolated from root sections and sheared into fragments between 200 and 300 bp in length. These fragments underwent end repair, dA tailing, and ligation to methylated adaptors for subsequent bisulfite conversion. Libraries were PCR amplified, purified, and quality controlled via Agilent DNA 1000 chip. Sequencing was done on a HiSeq 2500 v4 with 125 bp paired-end reads. Reads were trimmed and quality controlled using Trim Galore and FASTQC respectively (http://www.bioinformatics.babraham.ac.uk/projects/index.html). Reads were aligned and extracted with Bismark (Krueger and Andrews, 2011). Differential methylation quantification and visualizations were done with MethylKit (Akalin et al., 2012). Additional visualizations were performed with deepTools (Ramírez et al., 2016) after converting Bismark-aligned bam files to bigWig and then generating computeMatrix files for plotHeatmap and plotEnrichment outputs.

### Gene Regulatory Network Analysis

All available protein-protein interaction data was gathered from the STRING v11 database (Szklarczyk et al., 2018) for genes that have an affiliated K4me3 peak at their genic or promoter region, as annotated by Homer AnnotatePeaks. This phosphate responsive root gene regulatory network (GRN) was organized in a hierarchical radial layout in Cytoscape v3.8.2 to better visualize both transcription factor interactions and biosynthetic and metabolic pathways (Shannon et al., 2003). Gene nodes were then colorized for the RNA-seq log2 fold change from the lateral root region.

### Additional Analysis and Tools

DNA motif analysis for all sequencing data was performed by the MEME suite using the ANR method, minimum motif width of six, and an e-value threshold of less than or equal to 0.01. Assorted data handling and manipulation was performed with Samtools (Li et al., 2009), Bedtools (Quinlan and Hall, 2010), and UCSC software packages (Kent et al., 2010). Genome browser views were carried out with the Integrated Genome Viewer software (Robinson et al., 2011).

## Data Availability Statement

Sequencing data is available on the National Center for Biotechnology Information Sequence Read Archive (NCBI SRA: https://www.ncbi.nlm.nih.gov/sra). BioProject ID for sequencing files is PRJNA454504.

## Author Contributions

LK, JM, DW contributed to the experimental design. BH, JM, ZL, and LK grew the sorghum plants and collected the root tissue samples for library preparation. JM and MR generated the sequencing libraries. NG and XW performed bioinformatic and statistical analyses; NG wrote the manuscript. All authors contributed to the manuscript writing and editing process.

## Acknowledgements

We would like to thank Jon Shaff for their assistance in growing the sorghum plants and helping isolate root tissue for the RNA-seq and epigenetic assays. We would like to thank Robert Martienssen and the Martienssen laboratory for discussions and feedback on the epigenetic assays.

## Figure Legends

**Supplemental Figure 1. DNA-binding Motif Enrichment of CG Differentially Methylated Peaks**. MEME analysis of the 100 bp surrounding differential CG 5-methylcytosine modified regions in BTx623. All motifs were detected using the ANR method in the MEME suite program with minimum significance threshold of e < 0.01 or less.

**Supplemental Figure 2. DNA-binding Motif Enrichment of CHG Differentially Methylated Peaks**. MEME analysis of the 100 bp surrounding differential CHG 5-methylcytosine modified regions in BTx623. All motifs were detected using the ANR method in the MEME suite program with minimum significance threshold of e < 0.01 or less.

**Supplemental Figure 3. DNA-binding Motif Enrichment of CHH Differentially Methylated Peaks**. MEME analysis of the 100 bp surrounding differential CHH 5-methylcytosine modified regions in BTx623. All motifs were detected using the ANR method in the MEME suite program with minimum significance threshold of e < 0.01 or less.

**Supplemental Figure 4. K4/K27 Trimethylation Ratio Footprints Change from SP to LP**. Frequency histograms comparing the global H3K4me3/H3K4me3 counts ratios that cover genic, and 1000 bp upstream and 5000 bp upstream (starting from TSS) regions in the BTx623 genome. Upon LP, the count ratios tend to coalesce to a lower value, shrinking back from the longer tails of larger H3K4me3/H3K27me3 ratios in SP conditions. Only regions where K4me3 and K27me3 counts were greater than zero in both SP and LP were included in this analysis.

**Supplemental Figure 5. RNA-seq Expression Heatmaps of Selected Transcription Factor Family Genes**. Gene expression across root regions for **A**) WKRY, **B**) NAC, **C**) bZIP, **D**) B3-domain containing, and **E**) AP2 transcription factor family genes. GO enrichment of top-expressed transcription factors in LP in the lateral root regions returns auxin-activated signaling pathway (GO:0009734). The negatively expressed genes in LP in the apex region return response to chitin (GO:0010200) and abscisic acid-activated (ABA) signaling pathway (GO:0009738) as well as the auxin-activated signaling pathway (GO:0009734). Hierarchical clustering was performed with the built-in heatmap.2 function in rStudio.

**Supplemental Figure 6. Peak Occurrence of Selected Cysteine/Sulfur Metabolism Genes**. Integrated Genome Viewer display of H3K4 and H3K27 trimethylation enriched peaks on the promoter/genic region of SORBI_3008G179900 and SORBI_001G427300 gene models. The display shows the two biological replicates for each histone mark during LP (top four panels) and SP (bottom four panels). Only the K4me3 trimethylation peaks are prominent for these genes and the peak region becomes smaller in the LP condition for both replicates.

**Supplemental Figure 7. Model depicting DNA methylation and H3K4 and H3K27 trimethylation decreasing globally during growth under limiting phosphorus conditions**. This shift in epigenetic peaks results in more open chromatin for transcription factor binding, up-regulating the expression of certain transcription factors via a feedback loop and ultimately increasing the expression of several developmental, metabolic, and hormone signaling pathways in the lateral root apical region. This shift ultimately modulates the RSA to predominantly increase lateral root growth to better mine phosphorus from low P soils.

